# USP21 and OTUD3 Antagonize Regulatory Ribosomal Ubiquitylation and Ribosome-Associated Quality Control

**DOI:** 10.1101/407726

**Authors:** Danielle M. Garshott, Elayanambi Sundaramoorthy, Marilyn Leonard, Eric J. Bennett

## Abstract

Defects within mRNAs or nascent chains that halt ribosomal progression can trigger ribosome-associated quality control (RQC) pathways that facilitate mRNA and nascent polypeptide destruction as well as ribosome recycling. Failure to remove defective mRNAs or nascent chains can lead to the accumulation of cytotoxic protein aggregates and proteotoxic stress. We previously established that the E3 ligase ZNF598 catalyzes regulatory ribosomal ubiquitylation of specific 40S ribosomal proteins required for downstream RQC events. Utilizing an optical RQC reporter we identify OTUD3 and USP21 as deubiquitylating enzymes that antagonize ZNF598-mediated 40S ubiquitylation and facilitate ribosomal deubiquitylation following RQC activation. Overexpression of either USP21 or OTUD3 enhances readthrough of stall-inducing sequences as compared to knock-in cells lacking individual RRub sites suggesting that combinatorial ubiquitylation of RPS10 (eS10) and RPS20 (uS10) is required for optimal resolution of RQC events and that deubiquitylating enzymes can limit RQC activation.

## INTRODUCTION

Cellular protein homeostasis must continuously adapt to changing environmental conditions and exposure to extrinsic proteotoxic stressors that challenge cellular, tissue, and organismal health. A prominent source of proteotoxic stress arises during translation where transcriptional or mRNA processing errors can result in the translation of defective or truncated proteins and lead to the accumulation of toxic nascent protein products (Brandman and Hegde, 2016; Schuller and Green, 2018). Failure to remove these deleterious proteins can lead to aggregation which contribute to human pathologies including a wide range of neurodegenerative disorders (Gestwicki and Garza, 2012). A multitude of cellular quality control pathways have evolved to guard against the accumulation of these aberrant nascent polypeptides and maintain cellular homeostasis (Lykke-Andersen and Bennett, 2014). Both co-translational and ribosome-associated quality control (QC) mechanisms monitor protein biogenesis and initiate tightly coordinated responses to the production of defective protein products or other failures during translation. One such response, the ribosome-associated quality control (RQC) pathway, identifies actively elongating ribosomal complexes whose progression is halted due to a defect in the translating mRNA or emerging nascent chain (Brandman and Hegde, 2016). After the initial recognition events, the RQC response catalyzes the degradation of both the mRNA and nascent polypeptide, followed by ribosomal subunit recycling. It has been shown that RQC failures result in the production of aberrant protein products and an eventual accumulation of protein aggregates (Choe et al., 2016; Defenouillere et al., 2016; Yonashiro et al., 2016).

RQC pathways have been genetically well-characterized in *S. cerevisiae*, however much remains to be elucidated for the mammalian RQC pathway. These genetic studies in yeast along with biochemical characterization of both yeast and mammalian RQC factors have delineated a series of events that occur when ribosome progression is slowed enough to initiate a QC response (Joazeiro, 2017). In *S. cerevisiae*, the initial recognition event requires the ribosomal protein Asc1 and the ubiquitin ligase Hel2 (Brandman et al., 2012; Kuroha et al., 2010). Subsequent studies in mammalian cells revealed that the orthologous gene products, RACK1 and ZNF598, are required for proper resolution of ribosomes translating long poly(A) sequences (Juszkiewicz and Hegde, 2017; Sundaramoorthy et al., 2017). Characterization of ZNF598 and Hel2 revealed that these homologous ubiquitin ligases ubiquitylate conserved residues on 40S subunit proteins which are critical for the faithful execution of downstream RQC steps when ribosomes decode poly(A) sequences.

While it is clear that regulatory 40S ribosomal ubiquitylation (RRub) is required for downstream RQC events, the precise mechanistic role 40S ubiquitylation plays during RQC and what cellular factors serve as RRub regulators remain open questions. In yeast, Hel2 catalyzes ubiquitylation of RPS3 (uS3) and RPS20 (uS10) with K63-linked chains, which recruit factors that facilitate mRNA remodeling and degradation (Matsuo et al., 2017; Saito et al., 2015). Similarly, yet distinct, the mammalian ortholog ZNF598 catalyzes monoubiquitylation events on RPS10 (eS10) and RPS20 (uS10) that are required to initiate RQC events when decoding poly(A) sequences. Activation of the unfolded protein response in mammalian cells triggers a second set of RRub events on RPS3 (uS3) and RPS2 (uS5) that do not appear to function within the RQC pathway and whose function remains uncharacterized (Higgins et al., 2015). RPS2 (uS3) and RPS3 (uS5) RRub events were demonstrated to be reversible upon removal of UPR agonists indicating that RRub events, in general, may be regulated by deubiquitylating enzymes (Dubs). Here, we utilized an overexpression screen to identify two Dubs, USP21 and OTUD3, whose expression stimulates read-through of poly(A)-initiated RQC events. We demonstrate that USP21 and OTUD3 can directly antagonize ZNF598-mediated RPS10 (eS10) and RPS20 (uS10) ubiquitylation events. Further, we show that USP21 and OTUD3 expression results in augmented removal of ubiquitin from RPS10 (eS10) and RPS20 (uS10) following UV-induced RQC. Interestingly, overexpression of either OTUD3 or USP21 results in enhanced stall read-through compared to knock-in cell lines engineered to lack either RPS10 (eS10) or RPS20 (uS10) RRub sites indicating that combinatorial ribosomal ubiquitylation is required for optimal RQC function. Taken together, our results clearly establish that deubiquitylating enzymes can limit RQC activity.

## RESULTS

### Regulatory ribosomal ubiquitylation is reversible

Previous studies demonstrated that the regulatory ubiquitylation of RPS2 (uS5) and RPS3 (uS3) that is induced upon unfolded protein response (UPR) activation is reversed upon cessation of the UPR (Higgins et al., 2015). Based upon this finding, we reasoned that the ZNF598-catalyzed ubiquitylation of RPS20 (uS10) and RPS10 (eS10) that is critical for initiation of ribosome associated quality control (RQC) pathways may also be reversible. For clarity, we will refer to specific 40S ribosomal proteins by their historical names to avoid confusion between RPS10 (eS10) and RPS20 (uS10). To examine the reversibility of RRub events, we exposed 293T cells to UV radiation to stimulate RPS2, RPS3, RPS10 and RPS20 regulatory ubiquitylation and allowed cells to recover for increasing periods of time. Consistent with previous studies, UV exposure induced RPS2, RPS3, RPS10, and RPS20 regulatory ubiquitylation (Fig 1A) (Elia et al., 2015; Higgins et al., 2015). Interestingly, RPS10 and RPS20 ubiquitylation was induced 30 minutes following UV exposure whereas RPS2 and RPS3 ubiquitylation occurred 4 hours after UV treatment when RPS10 and RPS20 modification diminished (Fig 1A,B). RPS2 and RPS3 ubiquitylation similarly diminished from peak levels at 12 hours and returned to steady-state levels 24 hours after UV exposure (Fig 1A,B). RPS2 and RPS3 ubiquitylation occurred coincidently with eIF2*α* phosphorylation indicating that RPS2 and RPS3 RRub events occur in response to activation of the integrated stress response. This observation is consistent with previous studies demonstrating that RPS2 and RPS3 RRub events require eIF2*α* phosphorylation (Higgins et al., 2015). RPS2 and RPS3 RRub events are likely functionally distinct from the more immediate RPS10 and RPS20 ubiquitylation that may occur as a direct result of UV-induced ribosomal stalls. These results indicate that all tested RRub events are reversible.

**Figure 1.**
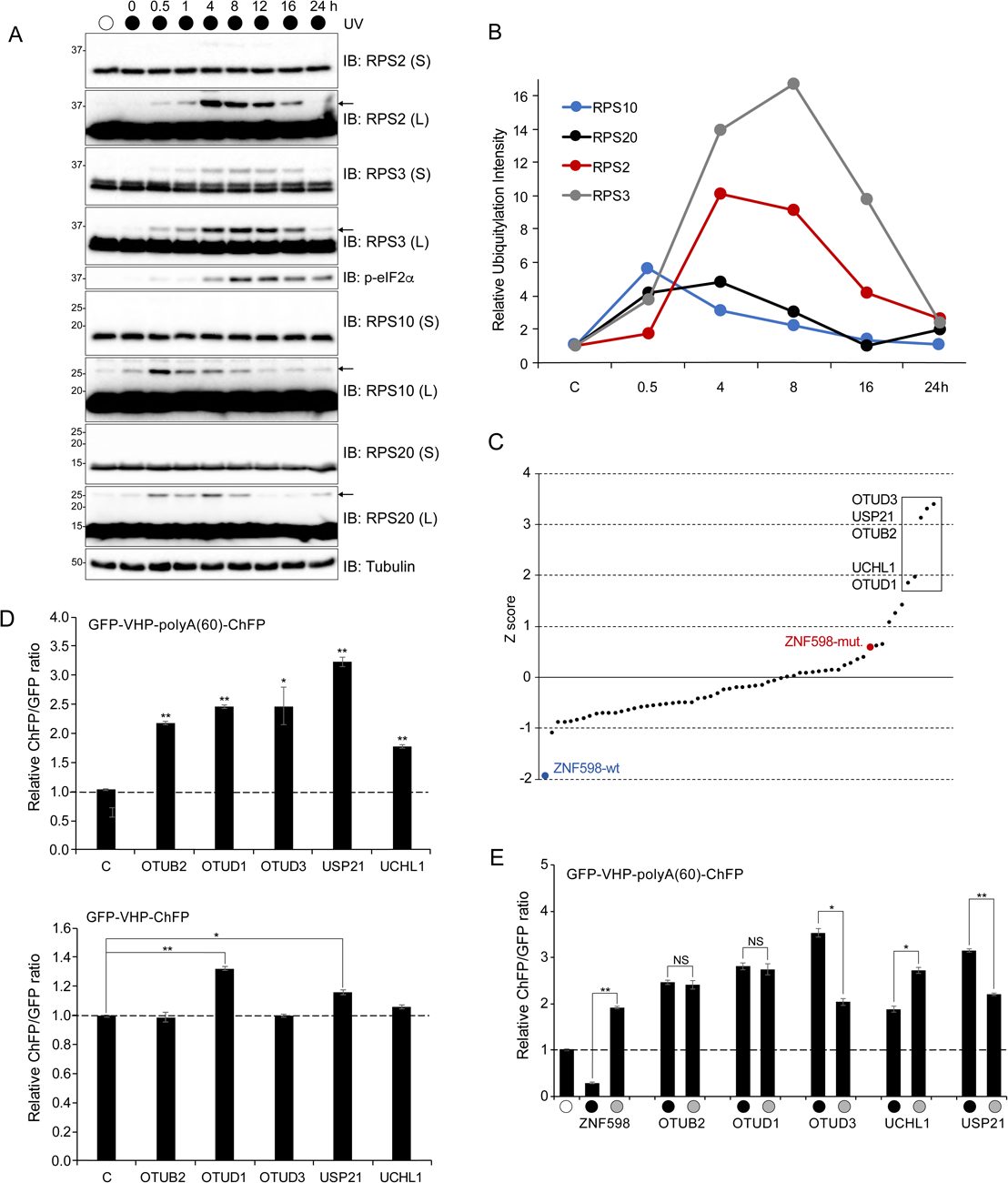
Identification of deubiquitylating enzymes that allow for readthrough of poly(A)-mediated RQC events. (A) 293T cells were exposed to UV and allowed to recover for the indicated times. Whole-cell extracts were analyzed by SDS-PAGE and immunoblotted with the indicated antibodies. The ubiquitin-modified 40S ribosomal protein is indicated by the arrow. S and L denote short and long exposures, respectively. (B) Quantification of the relative amount of ubiquitylated RPS10 (blue line), RPS20 (black line), RPS2 (red line) and RPS3 (grey line) during the UV treatment time course from the immunoblots in panel A. (C) A panel of 60 human deubiquitylating enzyme expression plasmids were transiently co-transfected with a dual-fluorescence polyA(60) reporter plasmid and the resulting ChFP and GFP fluorescence intensities were measured by flow cytometry. Depicted is the Z-score data representing the deviation of each individual ChFP:GFP ratio from the population mean ChFP:GFP ratio. Dubs with a Z-score greater than 1.5 are boxed to indicate candidate Dubs that induce increased ribosomal read-through of poly(A) stall sequences. Expression of wild type (blue dot) and catalytically inactive (red dot) ZNF598 are shown as controls. (D) 293T cells were co-transfected with either the poly(A)-containing reporter plasmid (top) or a reporter containing no stall sequence (bottom), along with expression plasmids for each of the candidate Dubs. The ChFP and GFP fluorescence intensities were measured by flow cytometry and the relative ChFP:GFP ratio is depicted for each Dub. Error bars denote SEM for triplicate transfections. **p<0.0001, *p<0.01 using Student’s t-test comparing Dub expression to the polyA(60) or no stall sequence control transfections. (E) 293T cells were co-transfected with individual expression plasmids for either wild type (black circle) or catalytically inactive (grey circle) candidate Dubs and the poly(A) reporter. Control transfections with the poly(A) reporter plasmid alone are indicated by the open circle. Fluorescence intensities were analyzed by flow cytometry and the relative ChFP:GFP ratio is plotted for each sample. Error bars denote SEM for triplicate transfections. **p<0.0001, *p<0.001 using Student’s t-test comparing the wild type to the catalytically inactive mutant for each Dub.

### Identification of deubiquitylating enzymes that antagonize RPS10 and RPS20 regulatory ubiquitylation

Our results implicate the direct involvement of deubiquitylating enzymes in regulating RQC function. To identify and characterize deubiquitylating enzymes (Dubs) that antagonize RRub events, we utilized a previously established dual-fluorescence reporter assay in which GFP and cherry fluorescent protein (ChFP) flank a villin-headpiece (VHP) linker sequence. Previous studies demonstrated that exogenous overexpression of this reporter plasmid in mammalian cells resulted in a liner correlation between GFP and ChFP protein levels (Juszkiewicz and Hegde, 2017; Sundaramoorthy et al., 2017). Introduction of a poly(A) sequence immediately after the VHP coding sequence reproducibly resulted in the repression of downstream ChFP fluorescence as compared to GFP, indicative of an RQC event initiating upon translation of the poly(A) sequence. Previous studies demonstrated that loss of ZNF598 function and the resulting decrease in RPS10 and RPS20 ubiquitylation results in readthrough of poly(A) sequences and a subsequent increase in the ChFP:GFP ratio of the stall reporter (Juszkiewicz and Hegde, 2017; Sundaramoorthy et al., 2017). Overexpression of a deubiquitylating enzyme that mediates deubiquitylation of RPS10 and RPS20 RRub events would be expected to phenocopy ZNF598 loss-of-function and enhance the amount of poly(A) readthrough. Based on this rationale, a panel of 60 human Dub expression plasmids were individually co-transfected with the poly(A) stall reporter plasmid into 293T cells and the corresponding ChFP:GFP ratio was measured by flow cytometry. Immunoblotting confirmed the expression of 58 Dubs (Fig. S1). A Z-score analysis of the ChFP:GFP ratio for the stall reporter identified five candidate Dubs whose expression resulted in enhanced poly(A) readthrough greater than 1.5 standard deviations above the population mean (Fig 1C). Confirmatory assays with the poly(A) stall-inducing reporter resulted in a reproducible enhancement of stall readthrough and a subsequent increase in the ChFP:GFP fluorescence ratio following overexpression of the five candidate Dubs (Fig 1D). To directly validate that the resulting increase in the ChFP:GFP ratio was specific to the poly(A) reporter, each candidate Dub was co-overexpressed with the control VHP plasmid without the internal poly(A) sequence (Fig 1D). OTUB2, OTUD3, USP10 and UCHL1 overexpression did not alter the ChFP:GFP ratio of the control reporter while OTUD1 and USP21 only modestly elevated the ChFP:GFP ratio indicating that the identified Dubs specifically alter the ability of ribosomes to progress through a poly(A)-induced ribosomal stall.

### Overexpression of candidate RQC-Dubs results in poly(A) stall-sequence readthrough in an activity-dependent manner

To examine whether the overexpression-induced increase in poly(A)-mediated stall readthrough was dependent on the catalytic activity of each of the identified Dubs, catalytically-inactive versions of each Dub were generated through site-directed mutagenesis to convert the critical catalytic cysteine residue to serine. Exogenous expression of each Dub and the respective catalytically-inactive mutant (CS) were co-expressed with the poly(A)-dual fluorescence reporter (Fig 1E). Overexpression of OTUB2 and OTUD1 catalytically-inactive mutants resulted in an equivalent degree of poly(A)-stall readthrough as compared to the respective wild type enzymes (Fig 1E). This result suggests that the observed increase in ChFP fluorescence does not require the catalytic activity of OTUB2 or OTUD1. Additionally, overexpression of the catalytically-inactive UCHL1 resulted in enhanced readthrough of the poly(A)-sequence compared to wild type UCHL1. In contrast, overexpression of the inactive mutants for the deubiquitylating enzymes USP21 and OTUD3 resulted in a substantial reduction of the ChFP:GFP ratios compared to wild type USP21 and OTUD3. Together, these results indicate that USP21 and OTUD3 expression results in elevated poly(A)-mediated stall readthrough in an activity-dependent manner.

### USP21 and OTUD3 antagonize ZNF598-mediated RRub events

Having demonstrated that exogenous expression of USP21 and OTUD3 enhanced poly(A) stall-induced readthrough, we wanted to examine the ability of the Dubs to directly antagonize ZNF598-mediated ubiquitylation events. The identified five candidate Dubs were transiently overexpressed alone or with exogenous ZNF598 and the poly(A) reporter. As expected, exogenous expression wild type, but not inactive ZNF598 resulted in decreased ChFP:GFP ratios as compared to control transfections (Fig 2A). Compared to UCHL1, OTUD1, and OTUB2 expression, both USP21 and OTUD3, when co-expressed with ZNF598, resulted in a greater than 2.5 fold increase in the ChFP:GFP poly(A) reporter ratio relative to what was observed when ZNF598 was expressed in isolation (Fig 2A). These results are consistent with the hypothesis that USP21 and OTUD3 directly antagonize ZNF598-mediated RRub events. To further examine the antagonism between ZNF598 and OTUD3 and USP21, parental HCT116 cells and ZNF598 knock-out (ZNF598-KO) cells were transiently co-transfected with either USP21 or OTUD3 expressing plasmids and the poly(A) stall-inducing reporter. We reasoned that overexpression of these Dubs in the absence of ZNF598 would not impact the amount of poly(A)-stall readthrough beyond that observed with loss of ZNF598 expression. As expected, overexpression of either Dub along with the poly(A)-reporter in parental HCT116 cells markedly increased the ChFP:GFP ratio of the stall reporter (Fig 2B,C). ZNF598-KO cells displayed the expected elevated ChFP:GFP ratio of the stall reporter which was only modestly enhanced upon further exogenous overexpression of either Dub (Fig 2B,C). These results demonstrate that the increase in the ChFP:GFP stall reporter ratio observed upon overexpression of OTUD3 or USP21 is highly dependent on the presence of ZNF598. The small increase in the ChFP:GFP ratio observed when overexpressing OTUD3 or USP21 in the ZNF598-KO cells suggests some possible ZNF598-independent role for Dub activity within the RQC pathway. Overall, these results demonstrate that USP21 and OTUD3 directly antagonize ZNF598.

**Figure 2.**
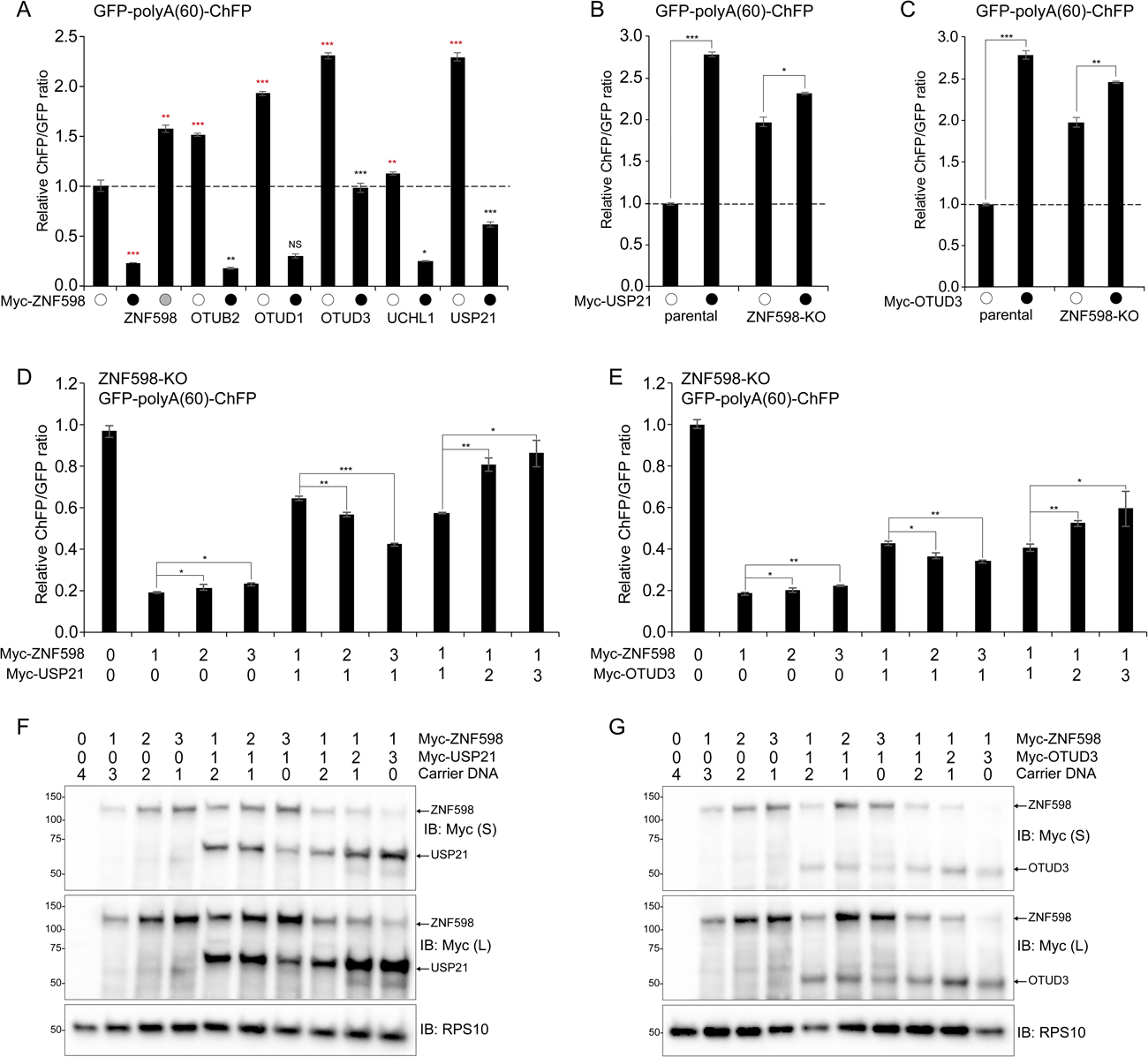
USP21 and OTUD3 directly antagonize ZNF598-mediated poly(A) readthrough in a dose-dependent manner. (A) 293T cells were transiently co-transfected with the poly(A) reporter plasmid and expression plasmids for each Dub alone (open circle) or with a wild type ZNF598 expression plasmid (black circle). Grey circle indicates expression of the catalytically inactive (C29A) ZNF598. The ChFP and GFP fluorescence intensities were measured by flow cytometry and the relative ChFP:GFP ratio is depicted. Error bars denote SEM for triplicate transfections. ***p<0.0001, **p<0.001, *p<0.05, using Student’s t-test comparing either the control transfection expression to samples expressing either ZNF598 or the indicated Dub alone (red asterisks), or comparing wild type ZNF598 expression alone to samples with co-expression of ZNF598 and the indicated Dub (black asterisk). (B-C) Parental HCT116 cells and ZNF598 knock-out (KO) cells were transiently co-transfected with USP21 (B) or OTUD3 (C) expression plasmids and the poly(A) reporter (black circles) or the poly(A) reporter alone (open circles). Fluorescence intensities were measured by flow cytometry and the relative ChFP:GFP ratio is depicted. Error bars denote SEM for triplicate transfections. ***p<0.0001, **p<0.001, *p<0.05, using Student’s t-test comparing Dub expression to the poly(A) reporter alone control transfection. (D-E) HCT116 ZNF598 knock-out (KO) cells were transiently co-transfected with increasing amounts of plasmid DNA for either wild type ZNF598 and USP21 (D) or OTUD3 (E) and the poly(A) reporter. Numbers indicate the ratio of transfected DNA for each plasmid. Error bars represent SEM of triplicate replicates. ***p<0.0001, **p<0.001, *p<0.05 using Student’s t-test comparing the different ZNF598 to Dub DNA ratios as indicated. (F-G) Whole-cell extracts from cells transfected as indicated in panels D and E were analyzed by SDS-PAGE and immunoblotted for the indicated antibodies. S and L denote short and long exposures, respectively.

### The abundance of ZNF598 in relation to USP21 or OTUD3 governs RQC events

Examination of quantitative proteomic datasets revealed that ZNF598 protein levels are 19-fold in excess of OTUD3 and USP21 levels were undetectable indicating that ZNF598 protein levels are in vast excess of either RRub Dub at steady state (Itzhak et al., 2016). Given the relative excess of ZNF598 compared to its antagonizing Dubs, we set out to examine how varying the levels of the Dubs relative to ZNF598 would impact RQC events. We transfected increasing amounts of a ZNF598 expressing plasmid in ZNF598-KO cell lines and examined poly(A)-mediated stall readthrough events using the stall reporter assay. Expression of the poly(A)-reporter with increasing concentrations of exogenous ZNF598 in isolation did not result in a dose-dependent decrease in the ChFP:GFP ratio suggesting that a low level of ZNF598 expression is required to fully restore RQC function and that elevated ZNF598 levels do not further enhance ribosomal stall resolution (Fig 2D-G). We then varied the relative ZNF598 expression levels compared to either USP21 or OTUD3 and examined the impact on the ChFP:GFP ratio of the stall reporter. When USP21 or OTUD3 and ZNF598 were expressed in an equimolar ratio, a substantial increase in the ChFP:GFP ratio was observed as compared to ZNF598 expressed alone which further verified the direct antagonism observed previously (Fig 2D,E). The reporter ChFP:GFP ratio declined as ZNF598 plasmid transfections where doubled and tripled with respect to USP21 or OTUD3 plasmid DNA (Fig 2D-G). Conversely, doubling and tripling the expression of USP21 while holding ZNF598 expression levels constant revealed additional readthrough of the poly(A) stall-inducing sequence with increasing ChFP:GFP ratios, suggesting further heightened antagonism of the ligase. This result suggests that maintaining ZNF598 expression levels high relative to USP21 and OTUD3 is a feature that may be required to enable rapid 40S ribosomal ubiquitylation upon RQC-triggering events that are not immediately removed by antagonistic Dubs.

### USP21 and OTUD3 catalyze deubiquitylation of regulatory RPS10 and RPS20 ubiquitylation events

Because overexpression of both OTUD3 and USP21 resulted in readthrough of the poly(A) stall-inducing reporter, we posited that both Dubs directly demodify ZNF598-catalyzed RRub events. To test this hypothesis, USP21 or OTUD3 were co-expressed with ZNF598 at expression levels three times greater than the ligase. Immunoblot analysis revealed that cells solely overexpressing ZNF598 displayed a 5-fold increase in the abundance of ubiquitylated RPS10 compared to untransfected cells (Fig 3A). Exogenous expression of USP21 substantially reduced the ZNF598-stimulated RPS10 ubiquitylation in an activity-dependent manner (Fig 3A). The same result was observed upon co-overexpression of OTUD3 and ZNF598 (Fig 3A). RPS20 ubiquitylation mirrored that of RPS10 in that the ZNF598-mediated enhancement of ubiquitylated RPS20 was reversed upon co-expression of either USP21 or OTUD3 in an activity-dependent manner. These results further demonstrate the ability of USP21 and OTUD3 to remove ubiquitin from RPS10 and RPS20 following ZNF598-mediated RRub events.

**Figure 3.**
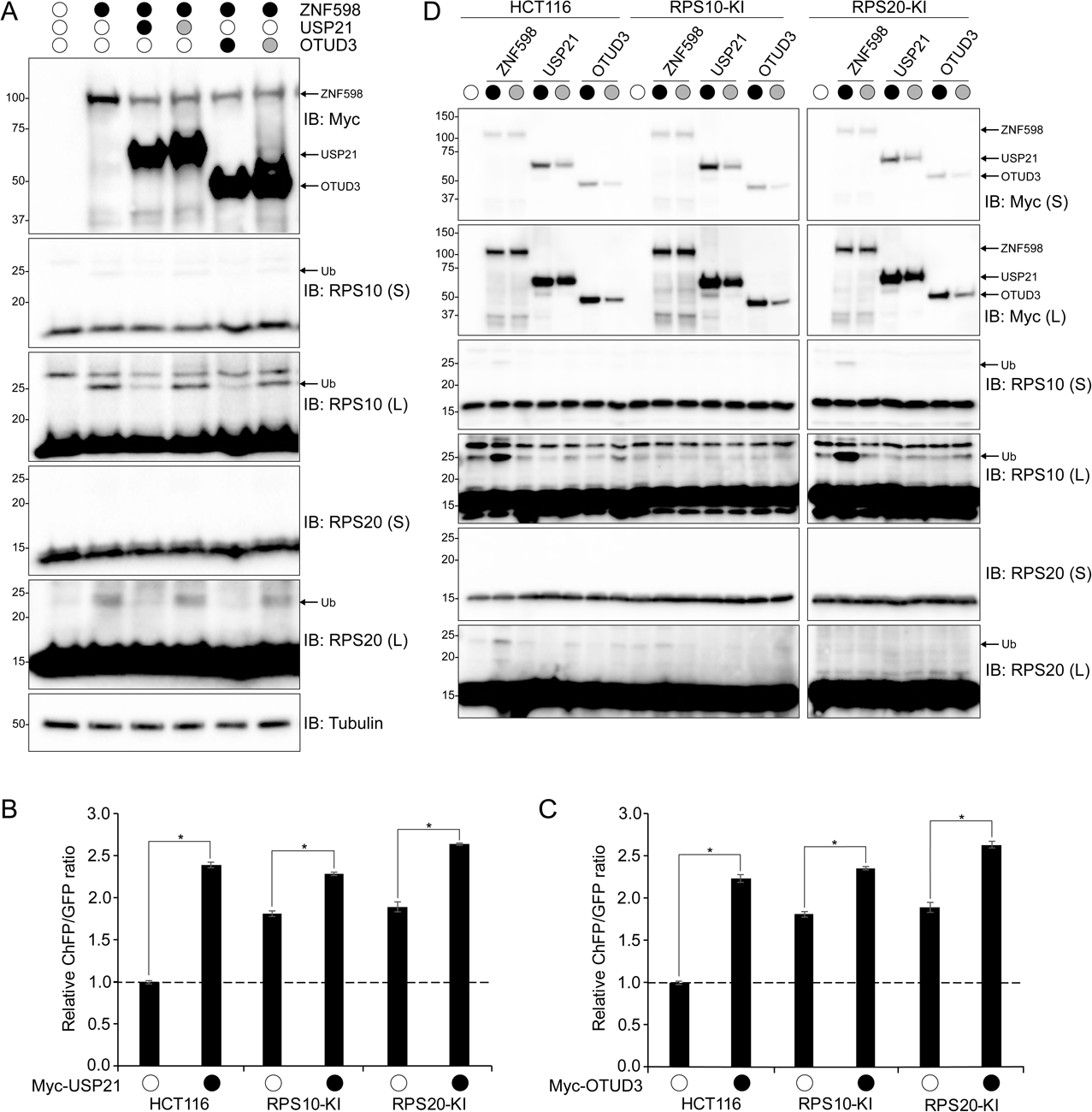
USP21 and OTUD3 deubiquitylate RRub events on RPS10 and RPS20. (A) HCT116 ZNF598-KO cells were co-transfected with expression plasmids for wild type (black circle) ZNF598, USP21, or OTUD3 and their respective catalytically-inactive mutants (grey circle). Whole-cell extracts were analyzed by SDS-PAGE and immunoblotted for the indicated antibodies. The ubiquitin-modified 40S ribosomal protein is indicated by the arrow. S and L denote short and long exposures, respectively. (B-C) Parental HCT116 cells and point mutant knock-in cell lines for RPS10 (RPS10-KI, RPS10K138RK139R) or RPS20 (RPS20-KI, RPS20K4RK8R) were transfected with the poly(A) reporter alone (open circles) or co-transfected with expression plasmids for USP21 (B) or OTUD3 (C) and the poly(A) reporter (back circle). ChFP and GFP fluorescence intensities were analyzed by flow cytometry and the relative ChFP:GFP ratio is depicted. Error bars denote SEM for triplicate transfections. *p<0.0001, using Student’s t-test comparing the expression of the Dubs to expression of the poly(A) reporter alone. (D) Whole-cell extracts from the samples analyzed in panels B and C were analyzed by SDS-PAGE and immunoblotted for the indicated antibodies. Black and grey circles denote expression of wild type or catalytically inactive Dubs, or ZNF598, respectively. The ubiquitin-modified 40S ribosomal protein is indicated by the arrow. S and L denote short and long exposures respectively.

To investigate the role of individual RRub events during RQC, we generated point mutant knock-in HCT116 cell lines in which the endogenous RPS10 or RPS20 loci were modified by CRISPR/Cas9 approaches to replace ubiquitylated lysine residues with arginine. We validated that RPS10 K138R/K139R knock-in (RPS10-KI) and RPS20 K4R/K8R knock-in (RPS20-KI) cell lines lacked the corresponding ubiquitylation events by immunoblotting cell lysates from untreated or RRub stimulated UV treated cells (Fig S2A,B). RPS20-KI cell lines completely lack UV-induced RPS20 ubiquitylation without impacting RPS10, RPS2, or RPS3 UV-induced RRub events (Fig S2A). As expected, RPS10-KI cells did not display UV-induced RPS10 ubiquitylation compared to control cell lines. However, RSP10-KI cells were also deficient in RPS20 UV-induced ubiquitylation while having a minimal impact on RPS2 and RPS3 ubiquitylation (Fig S2B). These results indicate that RPS10 ubiquitylation may be required for optimal RPS20 ubiquitylation upon induction of RQC events. Consistent with our previous results, RPS10-KI and RPS20-KI cell lines allowed for enhanced readthrough of poly(A)-mediated stall events compared to parental cells using our stall reporter FACS assay (Fig S2C) (Sundaramoorthy et al., 2017). To examine if the enhanced readthrough of poly(A) stall-inducing sequences observed upon USP21 or OTUD3 overexpression required RPS20 or RPS10 ubiquitylation, we expressed OTUD3 or USP21 in RPS20 or RPS10 knock-in cell lines along with the poly(A)-reporter. Expression of the poly(A)-reporter in both the RPS10-KI and RPS20-KI cell lines resulted in an elevated ChFP:GFP ratio as compared to the parental cells. USP21 or OTUD3 overexpression in either RPS10-KI or RPS20-KI cells resulted in a further enhancement of the ChFP:GFP ratio above the respective transfection controls. This result indicates that OTUD3 and USP21 can demodify both RPS20 and RPS10 and that this combined loss of RPS20 and RPS10 RRub events results in a stronger RQC defect compared to loss of either RPS20 or RPS10 individual ubiquitylation events. This is consistent with our immunoblotting data confirming that both USP21 and OTUD3 can remove ZNF598-mediated ubiquitylation of RPS10 and RPS20 (Fig 3A). To validate the poly(A)-reporter results, we immunoblotted cell lysates in which we overexpressed either wild type or catalytically inactive USP21 or OTUD3 in parental or RPS10-KI or RPS20-KI cell lines to visualize RPS10 and RPS20 ubiquitylation. As observed previously, overexpression of ZNF598 strongly stimulated ubiquitylation of RPS10 and RPS20 in parental HCT116 cells (Fig 3D). ZNF598 overexpression in RPS20-KI cells failed to induce RPS20 ubiquitylation without impacting the ability of ZNF598 to ubiquitylate RPS10. Conversely, ZNF598 overexpression in RPS10-KI cells could not ubiquitylate RPS10, and RPS20 ubiquitylation was substantially reduced compared to ZNF598 expression in parental cells. This is consistent with a model in which RPS10 ubiquitylation is needed prior to RPS20 ubiquitylation. While overexpression of wild type USP21 or OTUD3 reduced the abundance of both monoubiquitylated RPS10 and RPS20 in parental HCT116 cells, overexpression of the catalytically-inactive variants restored ubiquitylation to steady-state levels (Fig 3D). Overexpression of either Dub in the RPS10-KI cell line could further demodify the small amount of RPS20 ubiquitylation observed upon ZNF598 overexpression (Fig 3D). Similarly, USP21 and OTUD3 antagonized the ZNF598-dependent RPS10 ubiquitylation in RPS20-KI cells in an activity dependent manner. Taken together, these results indicate that USP21 or OTUD3 can deubiquitylate both RPS10 and RPS20 which results in enhanced readthrough of poly(A)-mediated stall events. Further, these results indicate that the combined ubiquitylation of RPS10 and RPS20 may be required for optimal resolution of RQC events.

### OTUD3 or USP21 loss-of-function has little impact on RQC and RRub

Having clearly demonstrated that OTUD3 and USP21 expression antagonizes ZNF598-mediated RQC events, we set out to test if OTUD3 or USP21 loss-of-function would result in enhanced resolution of poly(A)-mediated RQC events, mimicking ZNF598 overexpression. We knocked down OUTD3 or USP21 mRNA expression using three separate siRNA oligos and subsequently introduced the poly(A) stall reporter. Knockdown of USP21 or OTUD3 had no significant impact on the ChFP:GFP ratio compared to control siRNA transfections (Fig. S3A). ZNF598 overexpression resulted in the expected reduction in the ChFP:GFP stall reporter ratio indicating that OTUD3 or USP21 loss-of-function did not result in enhanced RQC function. We confirmed that the siRNA oligos substantially reduced endogenous OTUD3 protein levels and exogenous USP21 levels as USP21 antibodies failed to detect endogenous USP21 in these cell lines (Fig. S3B). Further, combined knockdown of USP21 and OTUD3 also failed to result in any significant reduction of the ChFP:GFP stall reporter ratio compared to ZNF598 overexpression (Fig. S3C). These results indicate that despite the robust and reproducible impact OTUD3 or USP21 overexpression has on RQC events, redundancy among endogenous Dubs precludes our ability to observe a loss-of-function phenotype for OTUD3 or USP21 on RQC function.

Because our Dub overexpression library did not include all known mammalian Dubs and the lack of a phenotype observed upon knockdown of USP21 or OTUD3 using the stall reporter assay, it remained plausible that a critical RQC Dub was overlooked. To examine if the Dubs not present in our overexpression library were possible RQC Dubs, we knocked down 24 Dubs and performed the poly(A) stall reporter assay. Examination of the ChFP:GFP ratio for the stall reporter revealed that knockdown of each of the 24 Dubs did not result in a significant reduction of the ChFP:GFP stall reporter ratio (Fig S3D). Only knock down of MINDY-2 (FAM63B) resulted in a small reduction of the ChFP:GFP ratio in our secondary screen. Given the K48-linkage preference observed for MINDY-2 and its reported nuclear localization, combined with the relatively small reduction in the ChFP:GFP ratio of the stall reporter compared to ZNF598 overexpression, we did not pursue MINDY-2 as a possible RQC Dub (Abdul Rehman et al., 2016; Kristariyanto et al., 2017). Collectively, these loss-of-function results indicate that no single Dub impacts RQC as robustly as ZNF598 and that possible redundancy in Dub function hinders our ability to observe a potent impact on our stall activity assay upon USP21 or OTUD3 loss-of-function.

### OTUD3 and USP21 specifically act to deubiquitylate 40S ribosomal proteins following RRub induction

To examine the ability of USP21 and OTUD3 to catalyze deubiquitylation of RRub events, we generated doxycycline (Dox)-inducible 293 cell lines that conditionally express either the wild type or catalytically-inactive mutant of each Dub. To observe the reversal of RRub events, we induced ribosome stalling and subsequent RRub using UV exposure. To test the impact of Dub overexpression, cells were either treated with or without doxycycline for 16 hours prior to UV exposure. Cells were then UV irradiated and allowed to recover for increasing periods of time. Based on our previously established reversibility of RRub events, we suspected that overexpression of wild type USP21 or OTUD3 would induce a more rapid removal of RPS10 and RPS20 ubiquitylation during recovery from UV-induced stress. Control cells without induction of Dub overexpression displayed induced RPS10 and RPS20 ubiquitylation immediately following UV treatment, followed by rapid demodification 4 hours after UV exposure (Fig 4A). As observed previously, RPS2 and RPS3 ubiquitylation was observed 4 hours post UV treatment which coincided with eIF2*α* phosphorylation indicating that RPS2 and RPS3 ubiquitylation occurs as part of the integrated stress response. RPS2 and RPS3 ubiquitylation declined 12 hours after UV exposure and eventually returned to steady-state levels by 24 hours despite the sustained levels of phosphorylated eIF2*α*. Interestingly, in cells overexpressing exogenous wild type USP21, the amount of detectable RPS10 ubiquitylation rapidly declined 1-hour post UV exposure, while RPS2 and RPS3 ubiquitylation was completely ablated (Fig 4A). These observations suggest that USP21 can demodify not only RPS10 and RPS20 following UV-induced stress, but also RPS2 and RPS3. To substantiate these observations, we induced the expression of catalytically-inactive USP21 and determined the dynamicity of UV-induced RRub events. Immunoblots confirmed that the dynamics of RPS2, RPS3, and RPS10 ubiquitylation following UV treatment in cells overexpressing catalytically-inactive USP21 were unaltered compared to cells without Dox-treatment. (Fig 4B). These findings confirm that USP21 can remove ubiquitin from RPS2, RPS3 and RPS10 in an activity-dependent manner. Similar to what was observed upon USP21 overexpression, OTUD3 expression resulted in substantially reduced RPS10 ubiquitylation following UV treatment compared to uninduced cells (Fig 4C). This reduction in RPS10 ubiquitylation was activity-dependent as induction of catalytically-inactive OTUD3 did not alter RPS10 ubiquitylation upon UV treatment (Fig 4D). In contrast to USP21, OTUD3 expression only marginally altered RPS2 and RPS3 ubiquitylation indicating that overexpression of OTUD3 impacts ribosome ubiquitylation in a less pervasive manner as compared to USP21 (Fig 4C,D). Interestingly, while overexpression of both OTUD3 and USP21 did not substantially reduce the levels of global protein ubiquitylation in cells, both USP21 and OTUD3 resulted in a reduction of the levels of ubiquitin-modified histones suggesting that overexpression of these Dubs can impact ubiquitylation of proteins beyond the ribosome (Fig 4A,C).

**Figure 4.**
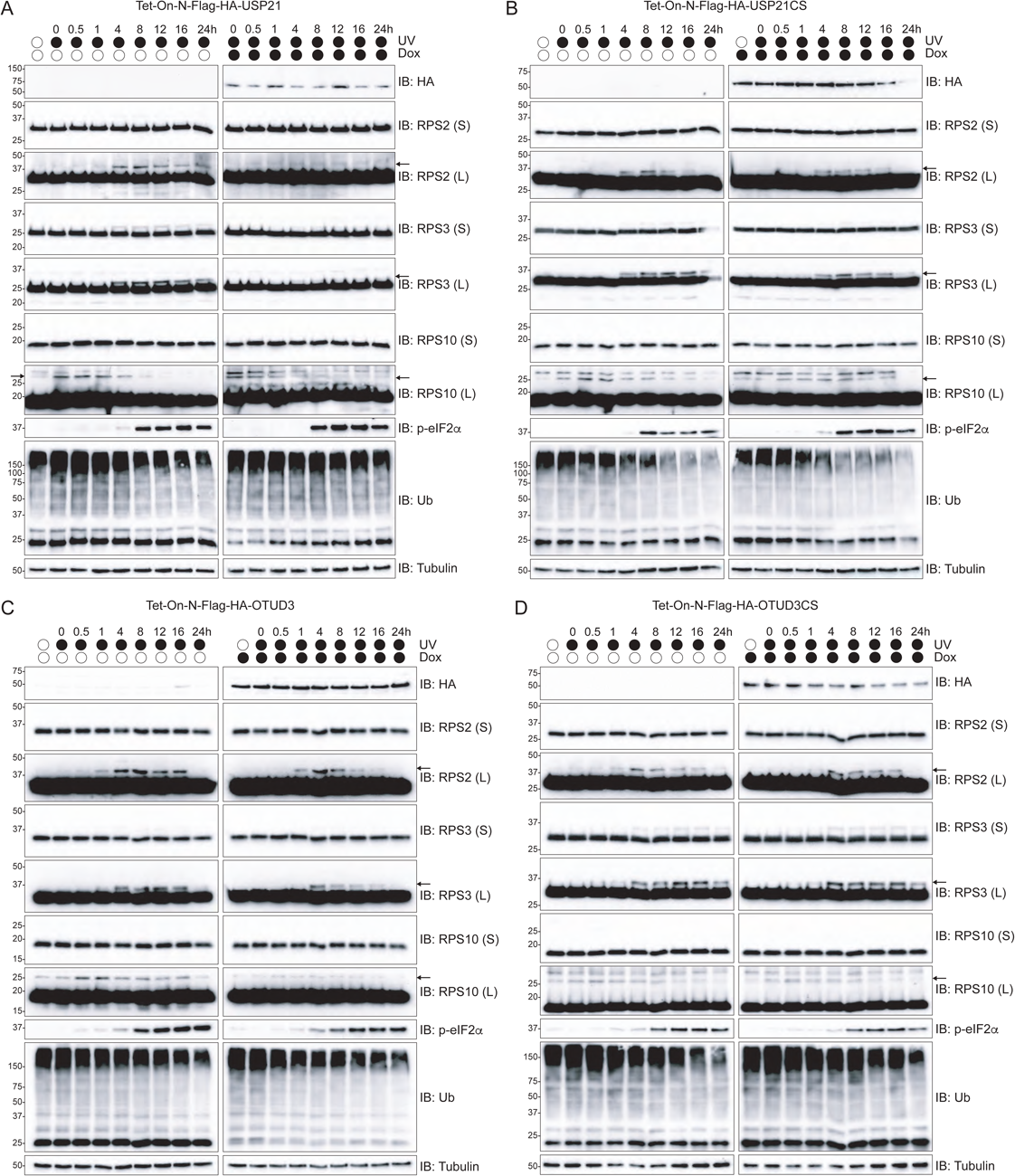
USP21 and OTUD3 expression accelerates RRub demodification following UV exposure. (A-D) Inducible 293 cells expressing either wild type USP21 (A), OTUD3 (C) or their catalytically inactive versions (B, D, respectively) in the presence (right panels) or absence (left panels) of doxycycline for 16 hours were exposed to UV, then allowed to recover for the indicated times. Cells were collected at the respective time points and whole-cell extracts were analyzed by SDS-PAGE and immunoblotted for the indicated antibodies. The ubiquitin-modified 40S ribosomal protein is indicated by the arrow. S and L denote short and long exposures, respectively.

To confirm the difference in the ability of USP21 and OTUD3 to impact RRub events, we again induced expression of wild type or catalytically-inactive USP21 or OTUD3 using our inducible cell lines. After induced Dub expression, cells were treated with dithiothreitol (DTT) or harringtonine (HTN) which stimulates RPS2 and RPS3 RRub events in a distinct manner (Higgins et al., 2015). Consistent with what we observed upon UV exposure, USP21 expression reduced both HTN and DTT-induced RPS2 and RPS3 ubiquitylation in an activity-dependent manner (Fig S4A). OTUD3 expression reduced RPS2 ubiquitylation, albeit to a lesser extent than UPS21 expression, and had no impact on HTN or DTT-induced RPS3 ubiquitylation (Fig S4B). Taken together with the results observed upon UV exposure, these results indicate that USP21 overexpression has a larger impact on all RRub events examined and that OTUD3 primarily demodifies ZNF598-catalyzed RPS10 and RPS20 ubiquitylation events.

Given the pervasive nature of USP21-mediated deubiquitylation of RRub events, we examined if overexpression of any catalytically active Dub would result in demodification of RRub events regardless of their impact on the stall-reporter assay. We constructed another set of inducible 293 cells with conditional expression of wild type or catalytically-inactive USP11. USP11 overexpression did not significantly alter the ChFP:GFP ratio of the stall reporter in our initial screen (Fig 1C). Overexpression of USP11 did not alter RPS2, RPS3, or RPS10 ubiquitylation dynamics following UV exposure (Fig S4C,D). This result indicates that, despite the promiscuous nature of USP21-mediated deubiquitylation of RRub events, general Dub overexpression is insufficient to antagonize RRub. Further, these results validate the ability of the initial screen to identify Dubs with the specific ability to demodify RRub events.

## DISCUSSION

Spatially distinct protein quality control (QC) mechanisms have been established within cells to respond to highly localized QC events. The endoplasmic reticulum-associated degradation (ERAD) pathway represents one of the most well-characterized, organelle specific quality control pathway that has evolved to respond to protein homeostasis stress occurring both within the ER lumen, and at the membrane interface with the cytoplasm (Ruggiano et al., 2014). The spatial separation allows for the utilization of specific QC factors that sense and execute QC functions for what can be distinct substrates. The ribosome represents another spatially distinct and unique quality control hub that monitors translational progression and nascent chain quality (Pechmann et al., 2013). A myriad of defects within the mRNA arising from transcriptional or mRNA processing errors can slow the progression of ribosomes such that a quality control mechanism is initiated (Schuller and Green, 2018). Activation and implementation of the RQC pathway is critical to identifying and rescuing ribosomes that have stalled on aberrant transcripts. Failure to recognize these ribosomes will ultimately result in continued translation of the defective mRNA and an inability to ubiquitylate and degrade the potentially proteotoxic nascent polypeptide. Previous studies have established that conserved site-specific regulatory 40S ubiquitylation (RRub) is among the first signaling events required for proper RQC execution (Garzia et al., 2017; Juszkiewicz and Hegde, 2017; Matsuo et al., 2017; Sundaramoorthy et al., 2017). While the ubiquitin ligase that catalyzes the RQC-specific RRub events has been characterized in both mammals and yeast, whether RRub demodification was a critical step in ultimate resolution of RQC events and the identity of potential RRub Dubs remained unknown.

Here we demonstrate both the reversibility of these conserved ubiquitylation events, and identify two deubiquitylating enzymes, USP21 and OTUD3, that directly antagonize ZNF598-mediated RRub events during RQC. We show that overexpression of these Dubs results in augmented removal of ubiquitin from 40S ribosomal proteins and enhanced readthrough of a poly(A) stall-inducing sequence. However, USP21 and OTUD3 have overlapping yet distinct ribosomal protein substrate specificities. The ubiquitin-specific proteases (USP) and the ovarian tumor (OTU) family make up the two largest Dub families. While USP Dubs are typically more promiscuous with regards to the types of polyubiquitin linkages they demodify (Faesen et al., 2011), OTU Dubs have been shown to exhibit ubiquitin-chain linkage specificity (Mevissen et al., 2013). Consistent with these observations, USP21 is the more promiscuous than OTUD3 in that USP21 can hydrolyze all four tested RRub events while OTUD3 preferentially demodifies ZNF598-catazlyed RPS10 and RPS20 ubiquitylation events. Despite the observed promiscuity of overexpressed USP21 to deubiquitylate all examined RRub events, the inability of USP11 to induce deubiquitylation of 40S ribosomal proteins establishes that overexpression of any deubiquitylating enzyme is not sufficient to antagonize RRub (Fig S4). These results clearly establish a regulatory role for Dubs within the RQC pathway.

Overexpression screens have long been employed to identify critical components for a variety of molecular pathways. However, results from such gain-of-function screens should be interpreted with caution, especially given the demonstrated promiscuity of deubiquitylating enzymes in vitro. Ideally, loss-of-function studies would validate the results observed upon overexpression of our candidate RQC dubs. We have previously shown that the extent of RPS10 and RPS20 ubiquitylation in cell extracts correlates with ZNF598 expression levels (Sundaramoorthy et al., 2017) and knockdown or knockout of ZNF598 results in a near ablation of RQC RRub events as well as clear phenotypes using the FACS-based poly(A) readthrough assay. Further, the enhanced poly(A) readthrough observed upon loss of ZNF598 function can be rectified by ectopic expression of small amounts of ZNF598. Taken together, the lack of an observable phenotype in the ribosome stall assay upon OTUD3 or USP21 knockdown suggests genetic redundancy in RQC Dub activity and that the overall levels of RRub at steady-state are mostly controlled by regulating ZNF598 activity or expression.

The factors that govern the regulation of these Dubs and the sequence with which the RRub modifications are removed is still unclear. We postulate two ways that Dubs may act as regulators of the RQC pathway. First, USP21 and OTUD3 may limit the activity of ZNF598 through direct antagonism to prevent spurious RRub. Unregulated 40S ubiquitylation could result in premature translational attenuation and destruction of properly processed mRNAs. Though plausible, the observed substochiometric ratio of OTUD3 and USP21 relative to ZNF598 suggests that Dubs may not directly limit ZNF598 activity. The low expression levels of USP21 and OTUD3 relative to ZNF598 may ensure that when progression of the ribosome is halted to a degree that requires RQC activity, the forward reaction is favored. A second possibility is that Dubs serve to strip the 40S of its ubiquitin modifications, following 80S splitting, to recycle 40S particles that are competent to reenter the translation cycle. Implicit in this model is that a ubiquitylated 40S prevents or modifies translation initiation events which has not been directly established. More in-depth biochemical studies are required to identify the factors and mechanism that regulate these reversible ribosomal regulatory ubiquitylation events, and the timing by which OTUD3 and USP21 exert their activity during RQC.

Proteomic based studies have revealed that a substantial amount of the ubiquitin-modified proteome may play a role in regulating cellular processes rather than targeting substrates for degradation (Kim et al., 2011). Several of these putative regulatory ubiquitylation events appear to be conserved, including many ribosomal ubiquitylation events. A key question that remains to be answered is: what is the function of these conserved regulatory ubiquitylation events? While it is clear that at least two of the regulatory ribosomal ubiquitylation events are needed for downstream RQC processes, the exact mechanistic role that ubiquitylation plays during the RQC is unknown. It is possible that ribosome ubiquitylation, particularly near the mRNA exit channel is required for the recruitment of mRNA decay and ribosome splitting factors necessary for downstream RQC events. As such, deubiquitylation of the 40S ribosome may be needed to remodel the exit channel to allow access to the defective mRNA and recruitment of RNA decay factors. Another possibility is that ribosome ubiquitylation represents a molecular timer that distinguishes ribosomes that are simply paused at a specific codon to allow for proper nascent chain folding or localization from those that are terminally stalled due to an unreconcilable defect in the mRNA. We have previously demonstrated loss of either RPS20 or RPS10 ubiquitylation results in a similar enhancement of poly(A) readthrough events. Here, we establish that antagonizing both RPS10 and RPS20 ubiquitylation through overexpression of USP21 or OTUD3 results in a further enhancement of poly(A) readthrough events. Furthermore, we demonstrate that loss of RPS10 ubiquitylation (RPS10-KI) results in a reduction in RPS20 modification suggesting that RPS10 is the first ubiquitylation event required for RQC initiation, and modification of RPS20 is partly dependent on RPS10 ubiquitylation. Taken together, these results suggest that the combined modification of both RPS10 and RPS20 is required for robust resolution of stalled ribosomes, and that these two modifications may not be occurring simultaneously, but rather in succession, implying a sort of ubiquitin code on the ribosome. Furthermore, it remains possible that the individual ubiquitylation events on RPS10 and RPS20 may not be taking place on the same ribosome, but rather occur on neighboring, collided ribosomes. This model then implicates ribosome collisions as the critical first event needed to induce the RQC pathway and suggests a situation, where upon collision with the trailing ribosome, ZNF598 mediates ubiquitylation of RPS10 on the leading stalled ribosome followed by ubiquitylation of RPS20 on the trailing ribosome. This could require a specific conformation of ZNF598 in order to traverse both ribosomes, or it is conceivable that following addition of the first ubiquitin the ligase moves upstream to the next site. Support for this model comes from previous studies in yeast that indicate ribosome collisions are critical events to initiate the no-go RNA decay pathway (Simms et al., 2017), and are required for upstream cleavage of the defective mRNA by an unknown endonuclease (Guydosh and Green, 2017). Having shown that both modification of RPS10 and RPS20 are required for a robust resolution of the stall it is probable that the collision with the leading ribosome triggers the first ubiquitylation events that signals recruitment of downstream RQC factors. Further biochemical analysis is needed to determine the exact mechanism and timing by which Dubs hydrolyze RRub, and the role of RRub events in recruitment of mRNA decay and splitting factors.

## EXPERIMENTAL METHODS

### Plasmids

The dual-fluorescence translation stall reporter plasmids were described previously (Sundaramoorthy et al., 2017). All Dub coding regions were cloned into Myc-tagging CMV expression vectors using Gateway cloning (Invitrogen). Site-directed mutagenesis was done utilizing PCR-based approaches, confirmed by sequencing and screened for expression by immunoblotting.

### Cell lines, transfections and siRNA

HCT116 and HEK293 cells were grown in DMEM (high glucose, pyruvate and L-Glutamine) containing 10% fetal bovine serum (FBS) and 1% penicillin/streptomycin and maintained in a 5% CO_2_ humidified incubator. Where indicated, cells were exposed to 0.02J/cm^2^ UV radiation using a Spectorlinker^TM^ XL-1000 (Spectronics) before harvesting or treated with 5mM DTT or 2 μg/mL Haringtonine for 4 hours before harvesting.

HCT116 knock-in cells (RPS20 K4R/K8R and RPS10 K138R/K139R) were generated using CRISPR/Cas9 genome engineering approaches by Biocytogen. Individual clones were first validated by genomic sequencing. Cells containing the desired point mutation were selected for validation by immunoblotting.

Stable doxycycline inducible cell lines expressing Flag-HA-tagged proteins were generated using the Flp-In^TM^ system (Thermo Fisher) through single locus integration, and Hygromycin selection. Flp-In T-Rex 293 cells were transfected with Flp-In expression vectors for gene of interest (listed in resource table) using TransIT 293 transfection reagent (Mirus) according to manufacturer guidelines. Cells seeded at 60% confluency were transfected for 24 hours followed by selection of stable expression clones with 100 μg/mL Hygromycin. Protein expression was induced with 2 μg/mL doxycycline 16 hours prior to UV exposure, drug treatment, or harvesting.

All cellular transient transfections were performed using Lipofectamine 2000 (Thermo Fisher) and siRNA knockdown transfections were performed using Lipofectamine RNAiMAX (Thermo Fisher) according to manufacturer guidelines.

### Dual-fluorescence translational stall reporter assay

All dual-fluorescent reporter plasmid cellular transfections were done using Lipofectamine 2000 according to manufacturer guidelines. Cellular ChFP and GFP fluorescence intensities were measured 48 hours following transfection on a BD LSRFortessa^TM^ X-20 cell analyzer (BD Biosciences). FACS data was analyzed using FlowJo (v10.4.1).

### Immunoblotting

Cell pellets were resuspended in denaturing lysis buffer (8 M urea, 50 mM Tris-Cl, pH 8.0, 75 mM NaCl, 1mM *β*-glycerophosphate, 1 mM NaF, 1 mM NaV, 40 mM NEM and EDTA-free protease inhibitor cocktail (Roche Diagnostics)) and kept on ice throughout preparation. Cell lysates were sonicated for 10s at output of 3 W with a membrane dismembrator model 100 (Fisher Scientific) with a microtip probe followed by centrifugation at 15,000 rpm at 4^°^C for 10 min. Supernatant protein concentrations were determined by BCA Protein assay (Thermo Fisher). Laemmli sample buffer with β-mercaptoethanol was added to cell lysates and heated for at 95^°^C for 10 min. Lysates were resolved on 12 or 15% SDS-PAGE gels then transferred to PVDF membranes (Bio-Rad) using Bjerrum semi-dry transfer buffer (48 mM Tris Base, 39 mM Glycine-free acid, 0.0375% SDS, 20% MeOH, pH 9.2) and a semi-dry transfer apparatus (Bio-Rad Turbo Transfer) at 25V for 30 min. Immunoblots were blocked with 5% blotting grade nonfat dry milk (APEX Bioresearch) in T-BST for 1 hour, followed by diluted primary antibody (listed in resource table) in 5% BSA over-night. Immunoblots were developed with Clarity Western ECL Substrate (Bio-Rad) and imaged on a Bio-Rad Chemi-Doc XRS+ system. Imagelab (Bio-Rad) software was used to process all blots with final images prepared in Adobe Illustrator.

## QUANTIFICATION AND STATISTICAL ANALYSIS

All FACS-based assays, plasmid transfections and siRNA transfections were done in triplicate (n = 3). The mean ChFP:GFP ratio and SEM were calculated and compared to K20-reporter transfection alone, parental cell type or control siRNA knockdown. Immunoblot quantification of the relative ubiquitin modification was calculated by normalization of the individual Ub intensities for each time point to that of the no UV control. Significance (p value) was calculated using an unpaired two-tailed Student’s t test using GraphPad Prism 7.0.

## AUTHOR CONTRIBUTIONS

Conceptualization, E.J.B and D.M.G; Methodology, E.J.B and D.M.G; Investigation, D.M.G, E.S, and M.L; Writing-Original Draft, D.M.G and E.J.B; Writing-Review & Editing, D.M.G, E.J.B, E.S, and M.L; Visualization, D.M.G and E.J.B; Funding Acquisition, E.J.B; Supervision, E.J.B.

## ACKNOWLEDGMENTS

We thank the Goldrath lab (UCSD) for providing assistance on all FACS-based experiments. This work was supported by a UCSD Cell and Molecular Genetics Training Program (T32GM007240) and a National Science Foundation Graduate Research Fellowship (DGE-1650112) (D.M.G), and the NIH (DP2-GM119132 and PGM085764) (E.J.B). E.S. is a fellow in the San Diego Center for Systems Biology (PGM085764).

## DECLARATION OF INTERESTS

The authors declare no competing interests.

## SUPPLEMENTAL FIGURE LEGENDS

**Figure S1.**
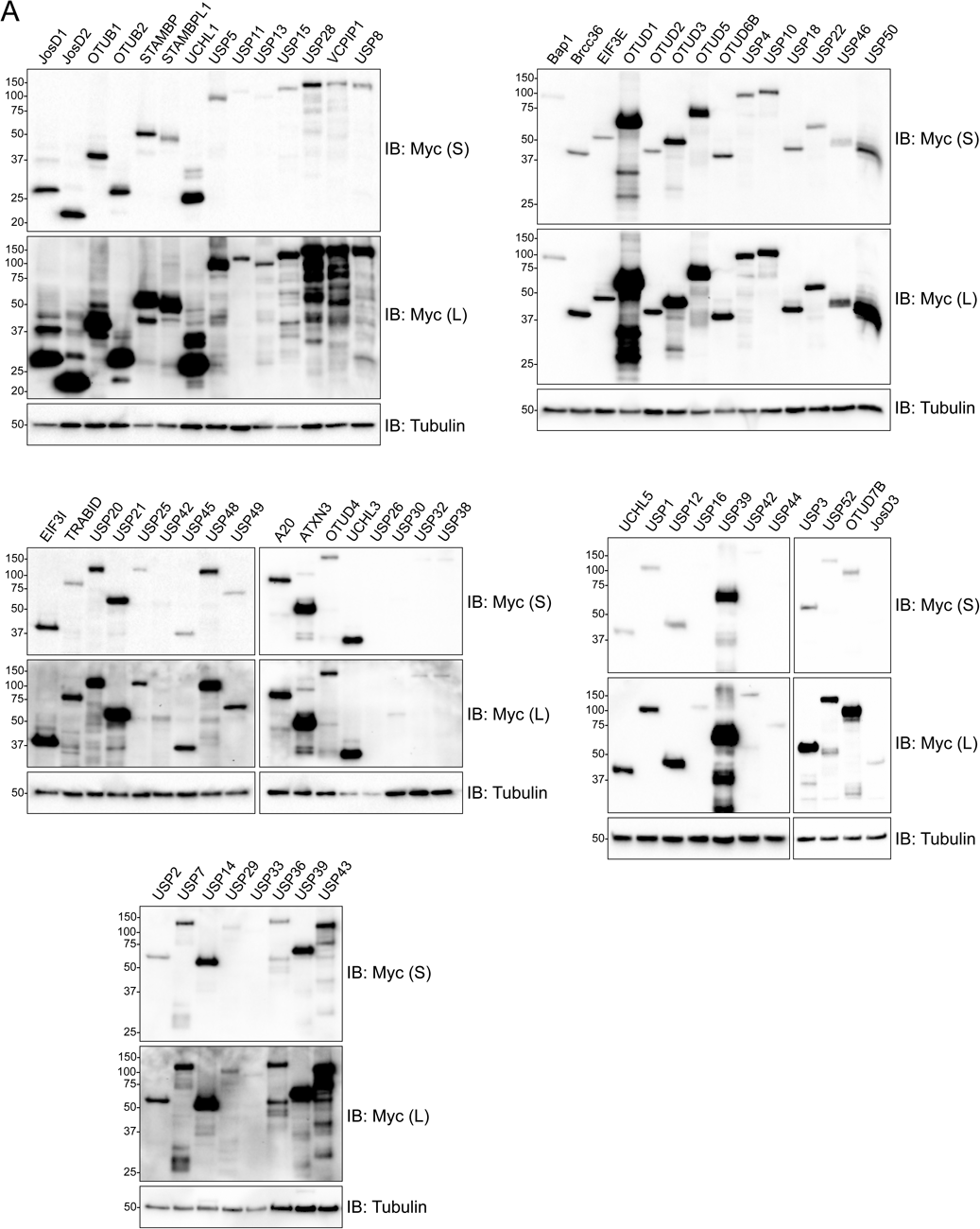
Validation of human Dub expression plasmids. (A) Whole-cell extracts from 293T cells transiently co-transfected with the indicated Myc-tagged Dub expression plasmid were analyzed by SDS-PAGE and immunoblotted with the indicated antibodies. S and L denote short and long exposures, respectively.

**Figure S2.**
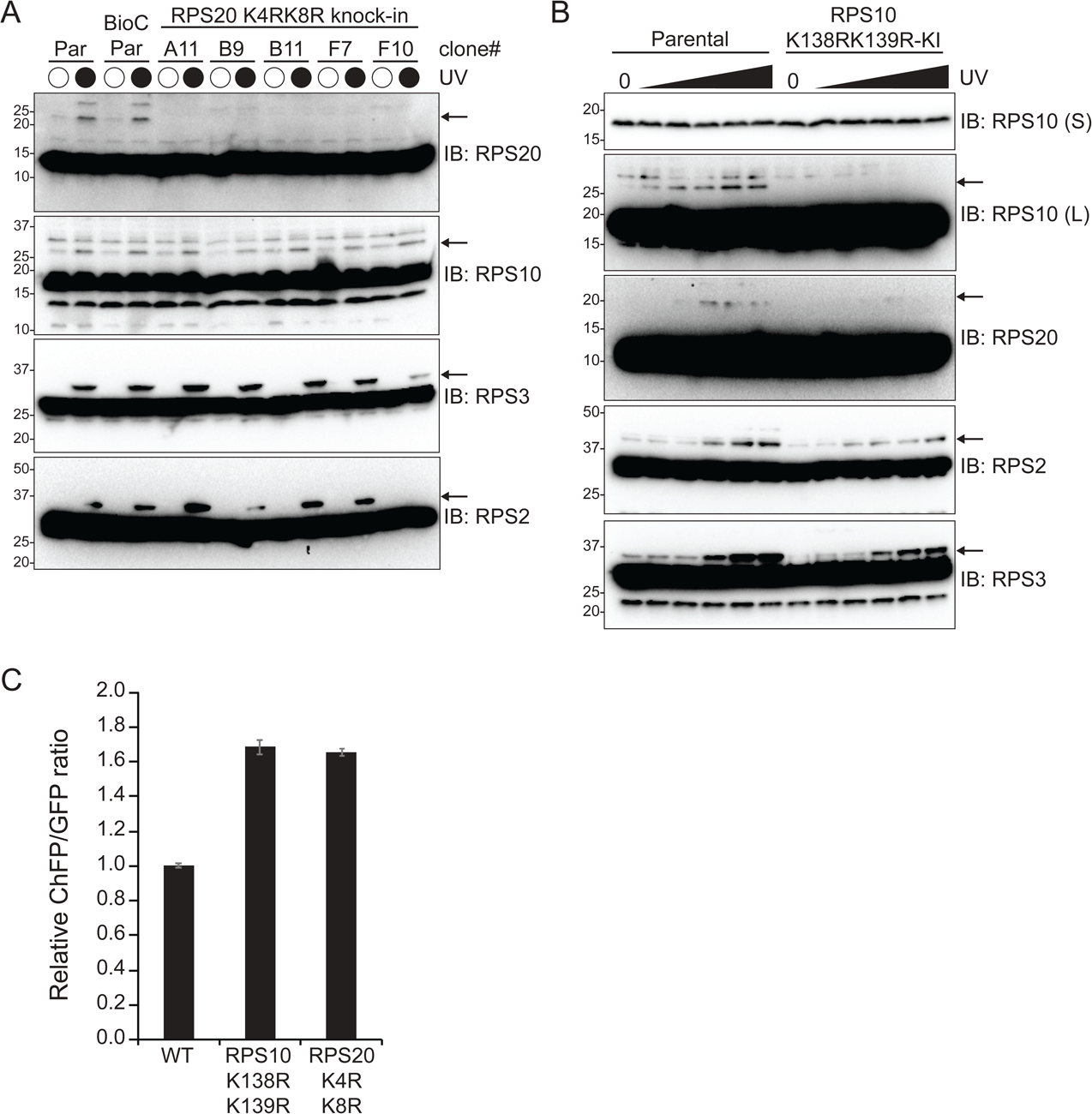
Validation of CRISPR/Cas9 engineered point mutant RRub knock-in cell lines. (A) Individual clones of HCT116 RPS20 knock-in (KI) cells where K4R and K8R point mutations were generated in the endogenous RPS20 locus or control parental (par) HCT116 cells were exposed to UV radiation to stimulate RRub events. Whole cell extracts were analyzed by SDS-PAGE and immunoblotted with the indicated antibodies. The ubiquitin-modified 40S ribosomal protein is indicated by the arrow. (B) HCT116 RPS10 knock-in cells where K138R and K139R point mutations were generated in the endogenous RPS10 locus or control parental (par) HCT116 cells were exposed to increasing doses of UV radiation. Whole cell extracts were analyzed by SDS-PAGE and immunoblotted with the indicated antibodies. The ubiquitin-modified 40S ribosomal protein is indicated by the arrow. S and L denote short and long exposures, respectively. (C) HCT116 RPS10-KI and RPS20-KI cells were transfected with the poly(A)-containing reporter plasmid. The ChFP and GFP fluorescence intensities were measured by flow cytometry and the relative ChFP:GFP ratio is depicted for each cell line. Error bars denote SEM for triplicate transfections.

**Figure S3.**
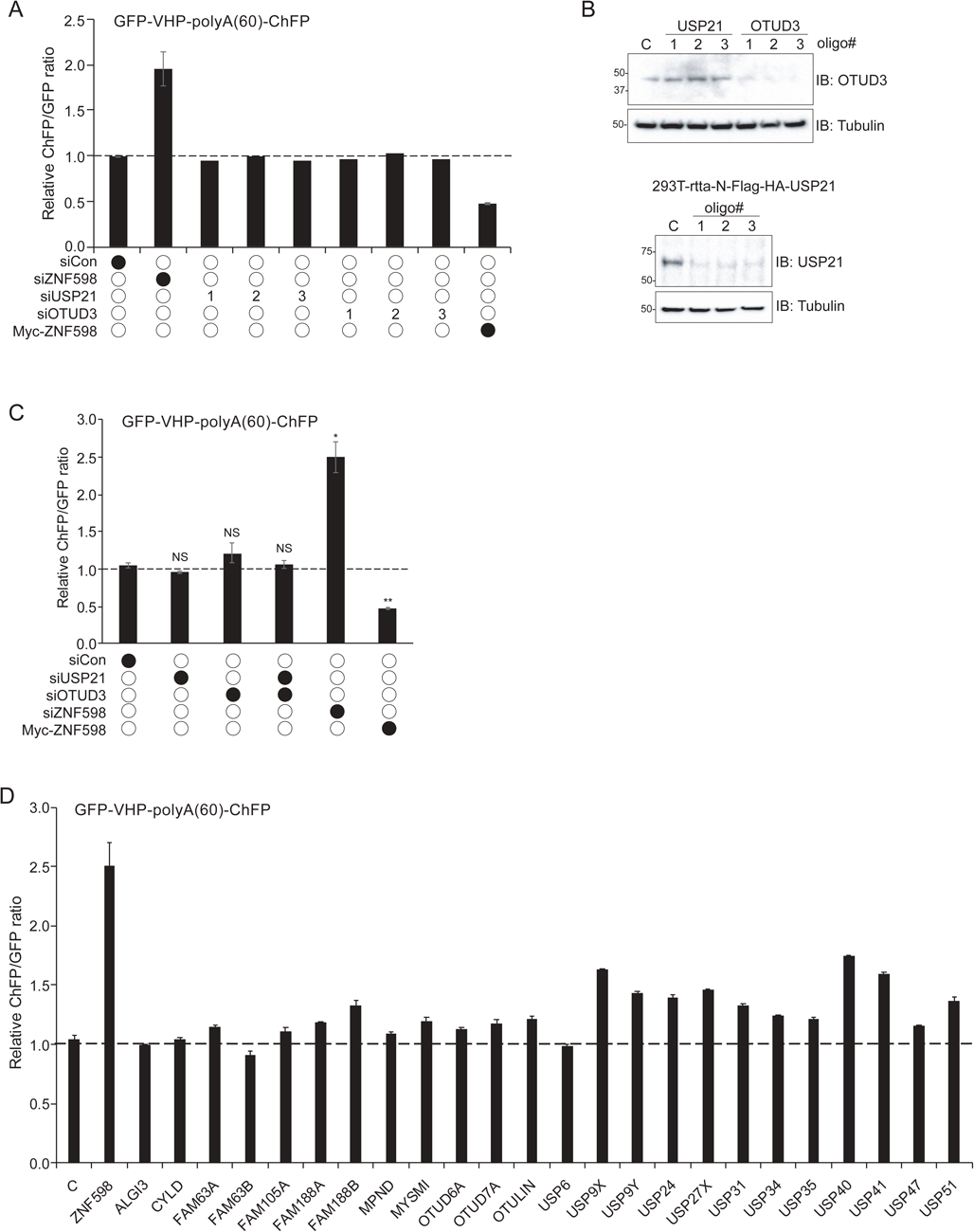
Knockdown of OTUD3 or USP21 does not result in enhanced resolution of poly(A)-induced RQC. (A) 293T cells were transfected with either control siRNA oligos or siRNA oligos targeting OTUD3, USP21 or ZNF598 individually. Numbers represent distinct siRNA oligos used to target OTUD3 or USP21. Cells were then transfected with the poly(A)-containing reporter and the ChFP and GFP fluorescence intensities were measured by flow cytometry. The resulting ChFP:GFP ratios are depicted. Error bars denote SEM for triplicate transfections. (B) Parental 293T (left) or inducible 293 cells (right) expressing exogenous Flag-HA-tagged USP21 were transfected with either control siRNA oligos or three separate siRNA oligos targeting USP21 or OTUD3. Cells expressing ectopic USP21 were induced with doxycycline 16h prior to cell collection. Whole-cell lysates were analyzed by SDS-PAGE and immunoblotted with the indicated antibodies. (C) 293T cells were transfected with control siRNA oligos or siRNA oligos targeting USP21, OTUD3, or ZNF598 as indicated. Cells were then transfected with the poly(A)-containing reporter and the ChFP and GFP fluorescence intensities were measured by flow cytometry and the relative ChFP:GFP ratio is depicted. Error bars denote SEM for biological replicates. **p<0.001, *p<0.01, using Student’s t-test comparing the different siRNA knockdowns or wild type ZNF598 overexpression to the siScramble control. (D) 293T cells were transfected with control siRNA oligos or pools of siRNA oligos (4 oligos each) targeting 24 individual Dubs or ZNF598. Cells were then transfected with the poly(A)-containing reporter and the ChFP and GFP fluorescence intensities were measured by flow cytometry. The resulting ChFP:GFP ratios are depicted. Error bars denote SEM for triplicate transfections.

**Figure S4.**
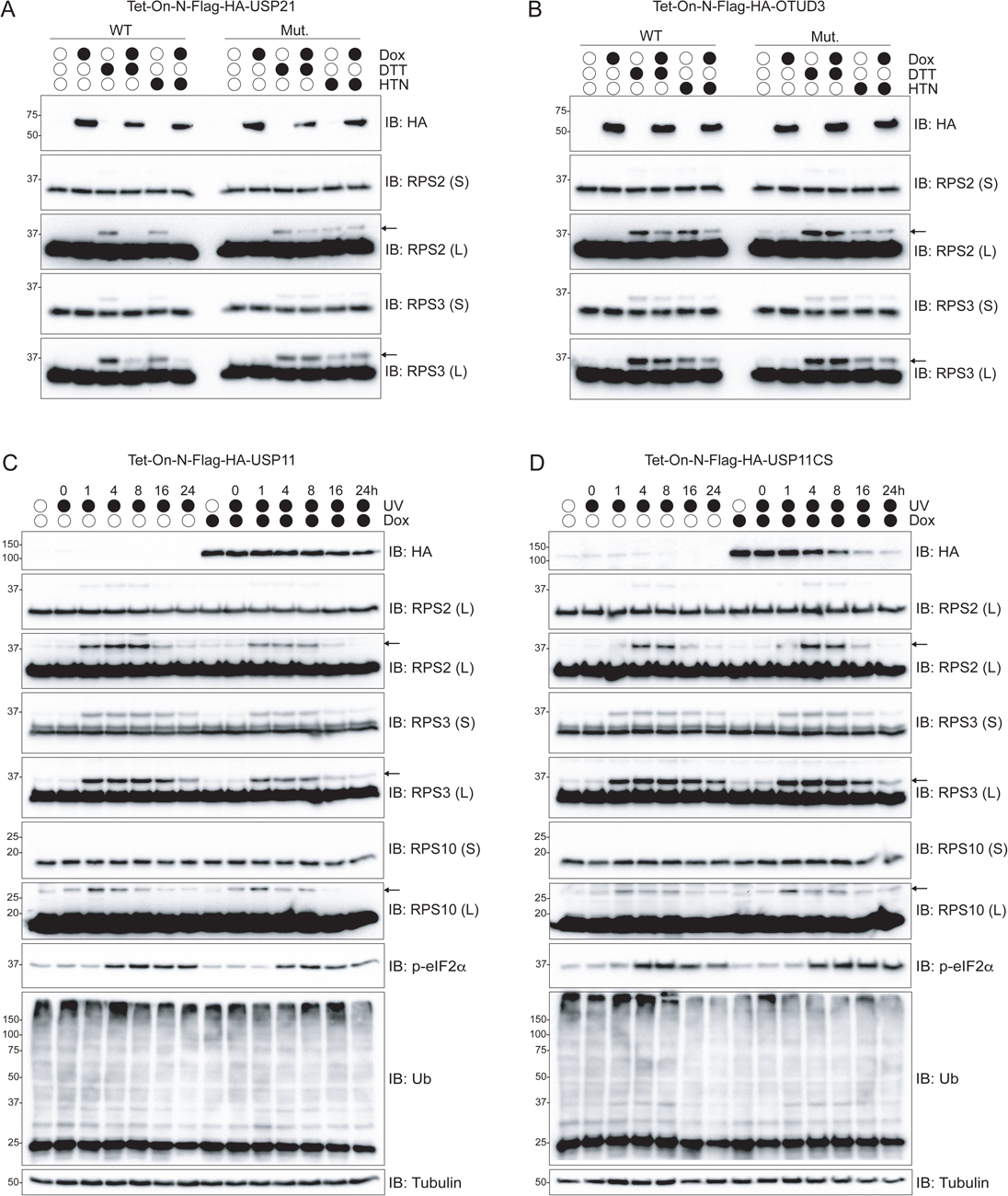
Examination of the site-specificity of RRub demodification upon Dub overexpression. (A-B) Inducible 293 cells expressing either wild type USP21 (A), or OTUD3 (B) or their catalytically inactive versions were induced with doxycycline for 16 hours and then treated with dithiothreitol (DTT) or harringtonine (HTN) to stimulate RPS2 and RPS3 RRub events. Cells were collected and whole-cell extracts were analyzed by SDS-PAGE and immunoblotted with the indicated antibodies. The ubiquitin-modified 40S ribosomal protein is indicated by the arrow. S and L denote short and long exposures, respectively. (C-D) Inducible 293 cells expressing either wild type (C) or the catalytically inactive USP11 (D) were either uninduced or induced with doxycycline for 16 hours as indicted. Cells were then exposed to UV and then allowed to recover for the indicated times. Cells were collected at the respective time points and whole-cell extracts were analyzed by SDS-PAGE and immunoblotted with the indicated antibodies. The ubiquitin-modified 40S ribosomal protein is indicated by the arrow. S and L denote short and long exposures, respectively.

## RESOURCE TABLE

## CONTACT FOR REAGENT AND RESOURCE SHARING

Requests for resources or further information can be directed to the Lead Contact Eric J. Bennett.

## SUPPLEMENTAL EXPERIMENTAL PROCEDURES

RESOURCE TABLE
| REAGENT or RESOURCE | SOURCE | IDENTIFIER |
| --- | --- | --- |
| **Antibodies** |  |  |
| Rabbit monoclonal anti-RPS2 | Bethyl Laboratories | Cat# A303-794A;<br>RRID:AB_11218192 |
| Rabbit monoclonal anti-RPS3 | Bethyl Laboratories | Cat# A303-840A;<br>RRID:AB_2615588 |
| Rabbit monoclonal anti-RPS10 | Abcam | Cat# ab151550 |
| Rabbit monoclonal anti-RPS20 | Abcam | Cat# ab133776 |
| Rabbit polyclonal anti-ZNF598 | Sigma-Aldrich | Cat# HPA041760;<br>RRID:AB_10792490 |
| Rabbit polyclonal anti-OTUD3 | Abcam | Cat# ab107646 |
| Rabbit polyclonal anti-USP21 | Abcam | Cat# ab171028 |
| Rabbit monoclonal anti-p-eIF2α | Cell Signaling Technology | Cat# 3398S |
| Mouse monoclonal anti-Ubiquitin | EMD Millipore/Chemicon | Cat# MAB1510;<br>RRID:AB_2180556 |
| Mouse monoclonal anti-HA | Biologend | Cat# MMS-101P;<br>RRID:AB_2314672 |
| Mouse monoclonal anti-tubulin | Cell Signaling Technology | Cat# 3873S;<br>RRID:AB_1904178 |
| Anti-Rabbit IgG (H+L), HRP Conjugate antibody | Promega | Cat# W4011;<br>RRID:AB_430833 |
| Anti-Mouse IgG (H+L), HRP Conjugate antibody | Promega | Cat# W4021;<br>RRID:AB_430834 |
| Lipofectamine RNAiMax | Thermo-Fisher | Cat# 13778030 |
| Lipofectamine 2000 | Thermo-Fisher | Cat# 11668019 |
| Mirus TransIT 293 | Mirus Bio llc | Cat# MIR 2700 |
| Protease inhibitor cocktail tablet | Roche | Cat# 11836170001 |
| Immun-Blot® PVDF Membrane | BioRad | Cat# 1620177 |
| Albumin Standard | Thermo Scientific | CAT# 23209 |
| Restore™ Western Blot Stripping Buffer | Thermo Scientific | Cat# 21059 |
| Clarity™ Western ECL Substrate | BioRad | Cat# 170-5061 |
| **Chemicals** |  |  |
| DL-Dithiothreitol | ACROS organics | Cat# 165680050 |
| Harringtonine | LKT labs | Cat# H0169 |
| Doxycyclin |  |  |
| Ammonium persulfate ($\geq 98\%$) | Sigma | Cat# A3678 |
| FBS | Mediatech (Corning) | N/A |
| Trypsin | Sigma-Aldrich | Cat# T1426 |
| NEM |  |  |
| Puromycin | Mediatech (Corning) | Cat# 61-385-RA |
| **Critical Commercial Assays** |  |  |
| BCA Protein Assay | Thermo Scientific<br>(Pierce) | Cat# 23225 |
| Experimental Models: Cell Lines |  |  |
| 293T | ATCC | Cat# CRL-3216 |
| HCT116 | ATCC | Cat# CCL-247 |
| 293T-FRT-FH-USP21-WT | This paper | N/A |
| 293T-FRT-FH-USP21-CS | This paper | N/A |
| 293T-FRT-FH-USP10-WT | This paper | N/A |
| 293T-FRT-FH-USP10-CS | This paper | N/A |
| 293T-FRT-FH-OTUD3-WT | This paper | N/A |
| 293T-FRT-FH-OTUD3-CS | This paper | N/A |
| 293T-FRT-FH-OTUD1-WT | This paper | N/A |
| 293T-FRT-FH-OTUD1-CS | This paper | N/A |
| 293T-FRT-FH-OTUB2-WT | This paper | N/A |
| 293T-FRT-FH-OTUB2-CS | This paper | N/A |
| 293Trex-iTAP-USP11_SL-WT | This paper | N/A |
| 293Trex-iTAP-USP11_SL-C275S | This paper | N/A |
| HCT116-ZNF598-KO | Sundaramoorthy et.<br>al. (2017) | N/A |
| HCT116-RPS10-KI | This paper | N/A |
| HCT116_RPS20-KI | This paper | N/A |
| Recombinant DNA |  |  |
| pCMV-GFP-2A-VHP-2A-CHFP<br>(linker control) | Juszkiewicz and<br>Hegde (2017) | N/A |
| pCMV-GFP-2A-VHP-K20-2A-CHFP<br>(K20 stall reporter) | Sundaramoorthy et.<br>al. (2017) | N/A |
| Sequence-Based Reagents |  |  |
| See Supplemental Experimental<br>Procedures for siRNA reagents |  |  |
| Software and Algorithms |  |  |
| FlowJo (v10.4.1) | BD biosciences |  |
| Prism 7 | GraphPad software |  |

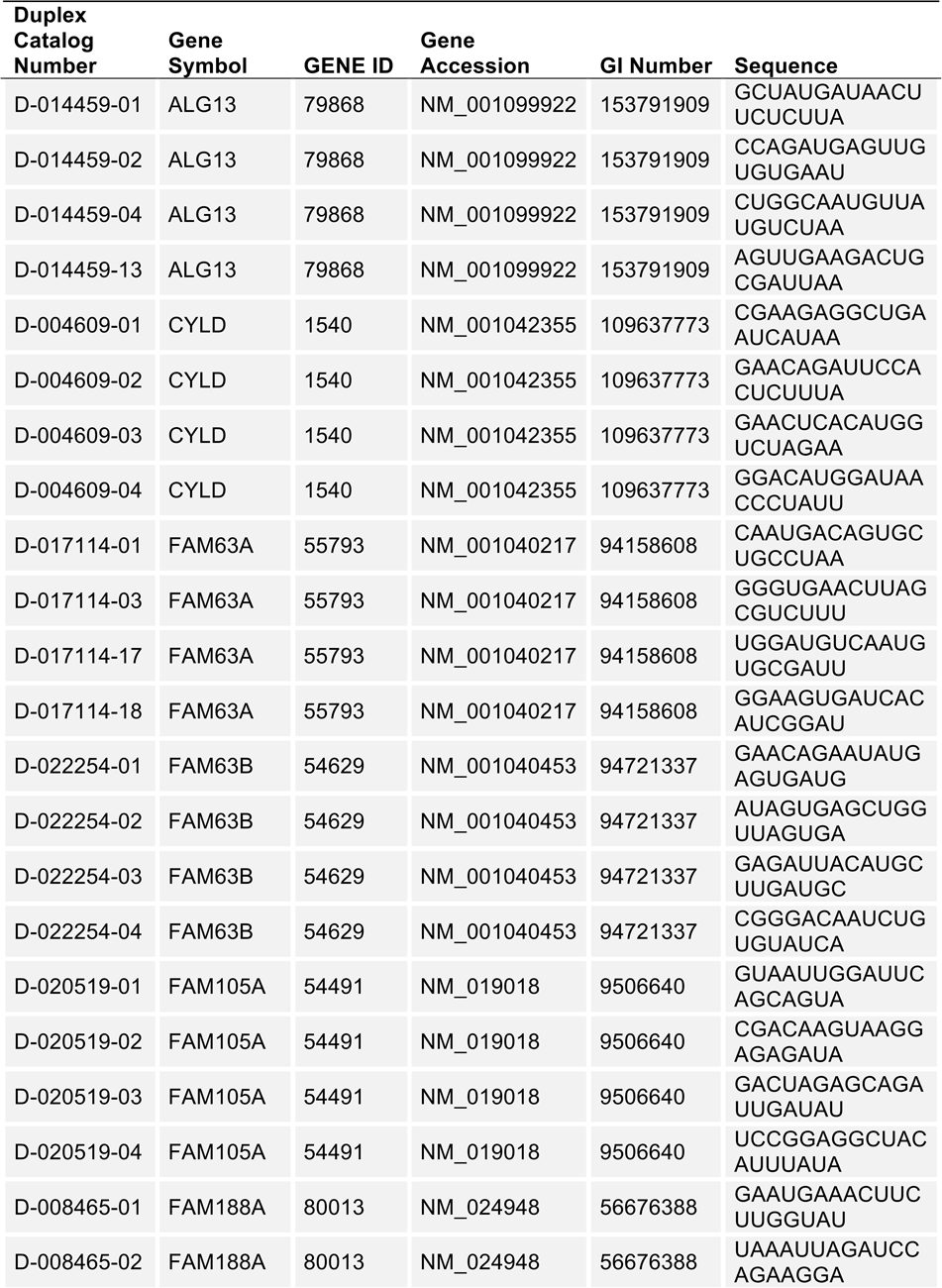
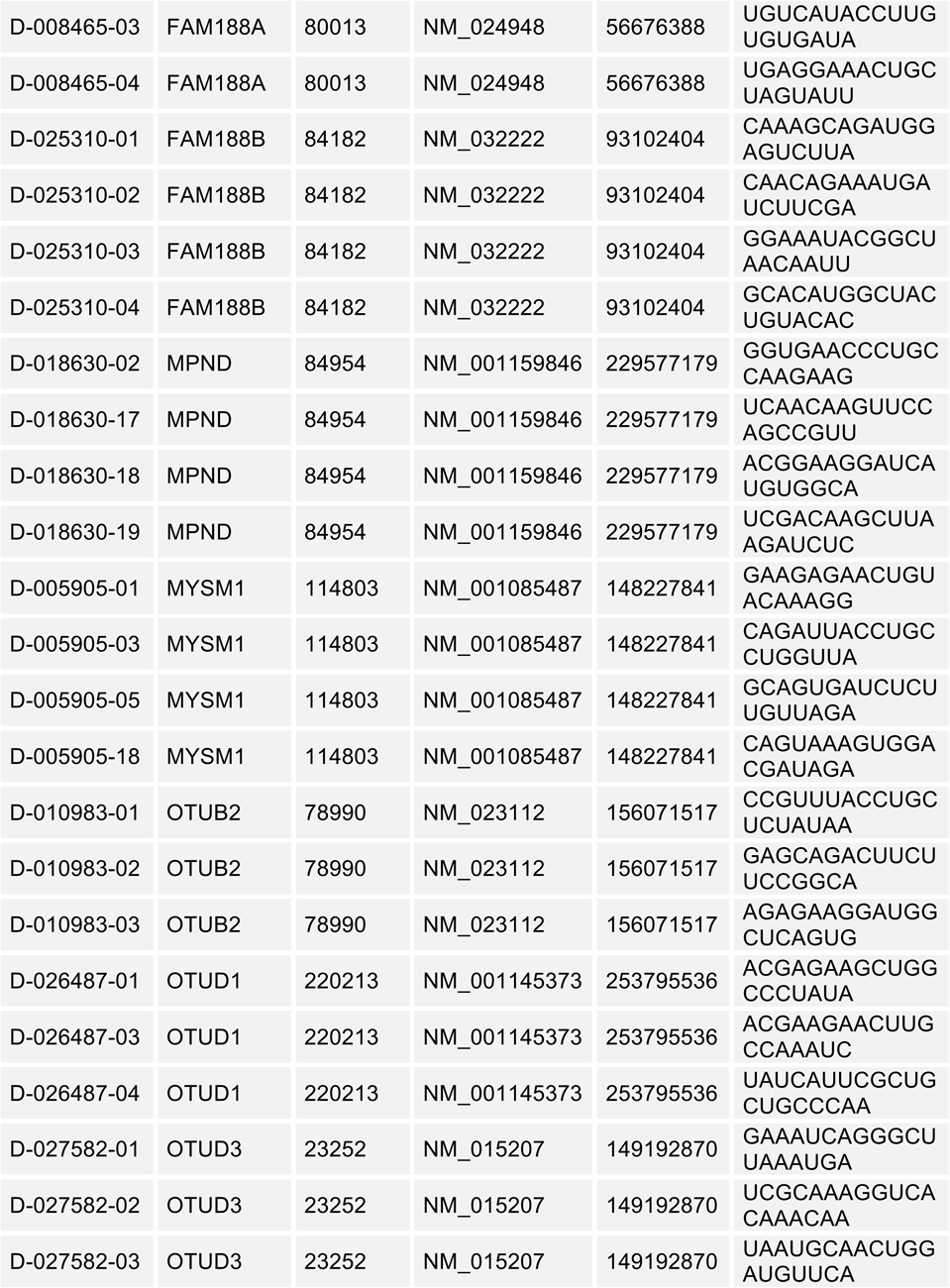
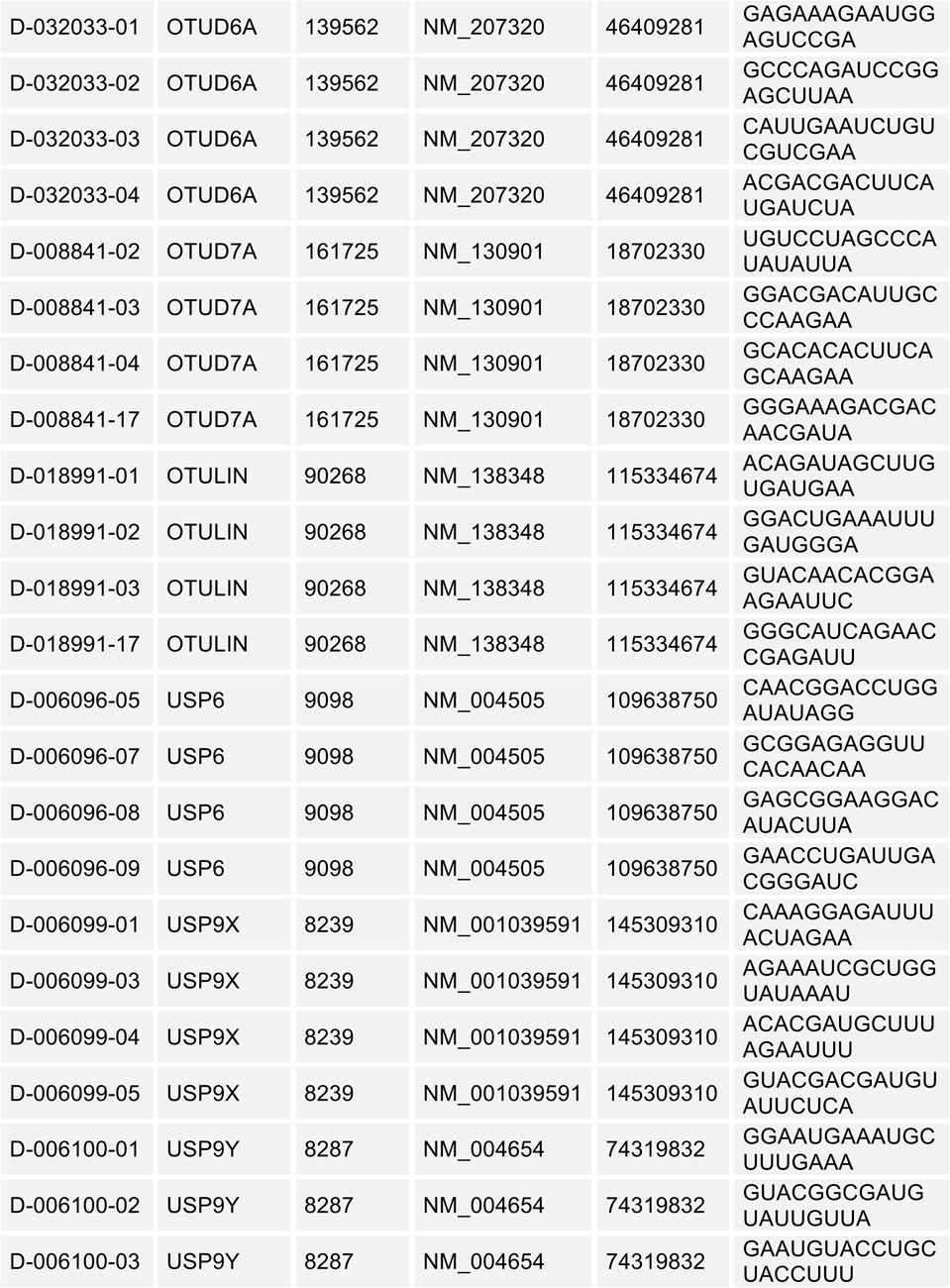
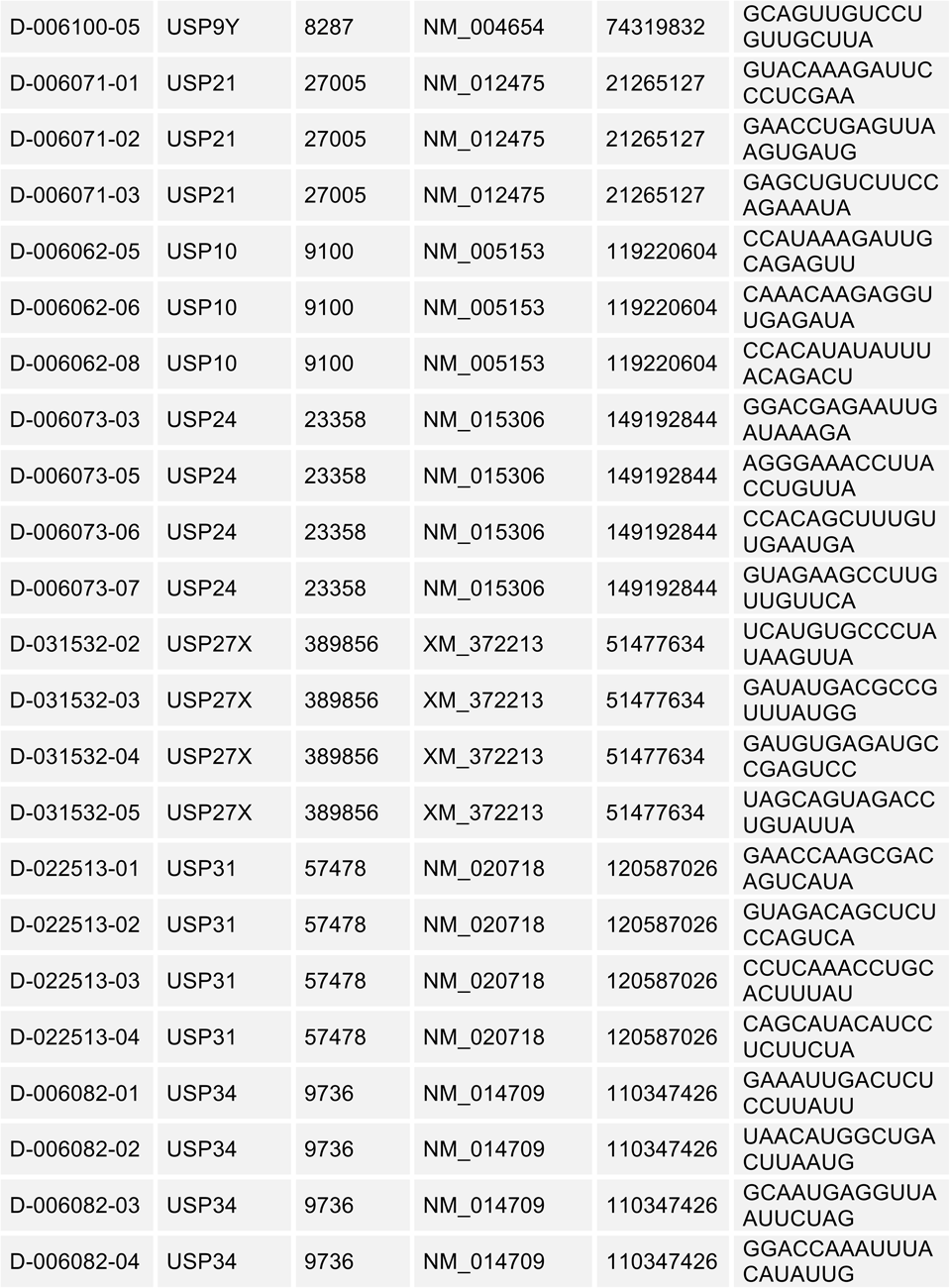
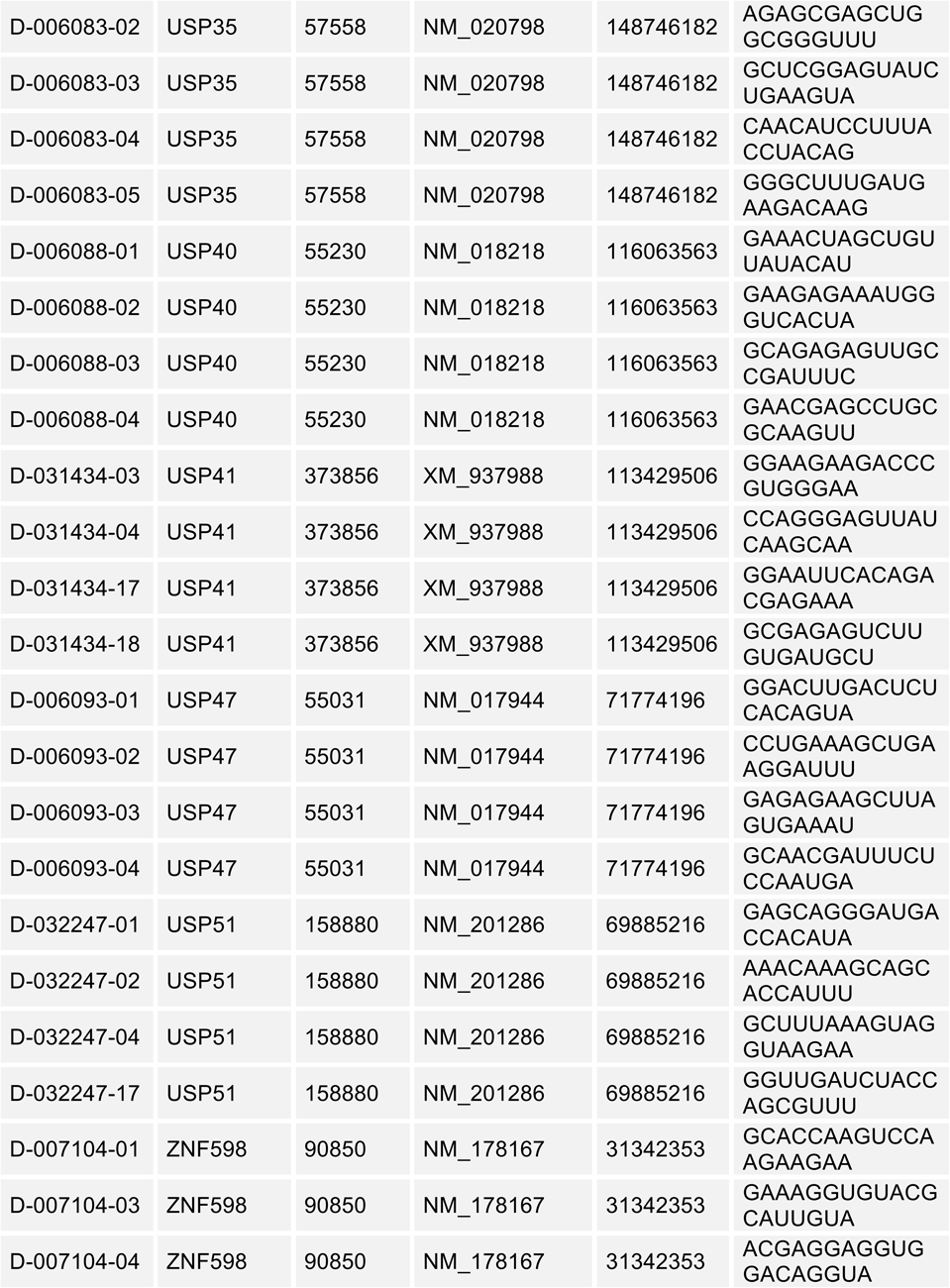
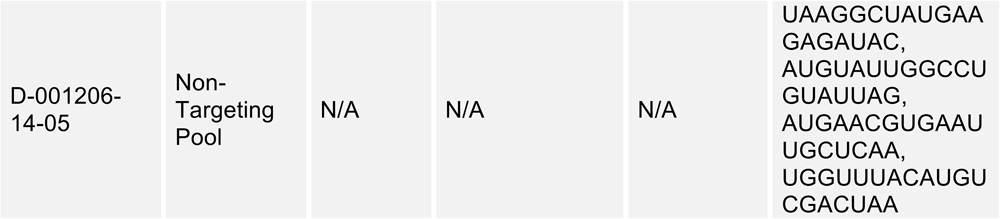
Table of RNAi oligonucleotides

